# High school science fair: Characterizing student communication and presentation practices and their association with SEF outcomes

**DOI:** 10.64898/2026.06.10.731420

**Authors:** Frederick Grinnell, Simon Dalley, Joan Reisch

**Author notes:** Corresponding author: E-mail address (FG).

## Abstract

*Obtaining, Evaluating, and Communicating Information* is one of the eight Science and Engineering Practices in the National Research Council framework that underlies Next Generation Science Standards (NGSS). Scientific inquiry requires evaluation and communication of research findings. These skills are part of scientific inquiry itself. The goal of this study was to characterize the communication practices students used during Science and Engineering Fairs (SEFs). To examine these practices, we included the following question in our online anonymous and voluntary national high school SEF surveys carried out during 2021-22 and 2022-23, “What types of communication and presentation skills did you use in your science fair project?” The possible answers were *literature review; research notebook; software to prepare tables, graphs, or images; written report; poster board preparation; PowerPoint presentation; and interview with the judges*. Literature review and research notebook are part of developing research questions and data collection. Software to prepare tables, graphs, or images, and PowerPoint presentation contribute to analyzing research and developing a presentation. Written reports, poster board preparation, and interviews with the judges provide the opportunity to present the findings. Overall, 1789 students answered the question. The percentage of students utilizing these skills ranged from 17.3% doing a literature review to 67.6% preparing a poster board. On average, students indicated the use of three skills. Poster board preparation and interview with the judges were selected by students more than twice as frequently as literature review and research notebook. More positive SEF outcomes were associated with a greater number of skills used by students. Among the different skills, use of a research notebook showed the strongest association. In this paper, we present the findings and discuss implications regarding use of notebooks and other communication and presentation skills in students’ SEF experiences and engagement with scientific inquiry.

## Introduction

Next Generation Science Standards (NGSS) emphasizes student experience of the practices of science as one of three essential dimensions of science education -- *students cannot comprehend scientific practices, nor fully appreciate the nature of scientific knowledge itself, without directly experiencing those practices for themselves* [1, 2]. How best to accomplish hands-on practice of science and engineering remains a work in progress [3-5]. Science and engineering fairs (SEFs) provide one important opportunity for hands-on practice. Begun almost one hundred years ago, the stated goals were the same as contemporary STEM education -- *to aid in the development of the scientific leaders of the next generation and at the same time foster a better understanding of science among its laymen* [6]. Now, SEFs have become part of science education not only in the United States, but also worldwide [7-16]. At the highest level, approximately 170,000 students from 400 science fairs compete to participate in the International Science and Engineering Fair (ISEF) [17].

SEFs potentially can promote three important and desirable STEM outcomes: (i) mastery of science and engineering (S&E) practices; (ii) interest in STEM; and (iii) interest in STEM careers [7, 11, 18, 19]. The idea that SEF participation could have a positive impact on high school students is consistent with research showing that science project-based learning advances students’ STEM understanding and interests at both the high school [20-25] and undergraduate levels [26-29]. Also, innovative high school programs that combine student participation in SEFs with student and teacher support promote STEM engagement and learning for all students including those from under-represented ethnic minorities and low socioeconomic backgrounds [30-35]. In general, student engagement in STEM research projects including SEFs fits into the self-determination model in education [19, 36-38], which focuses on the importance of student autonomy, competence, and community engagement [39, 40].

The goal of our previous research has been to describe and characterize students’ experiences in high school SEFs. To accomplish this goal we developed a mixed methods approach using anonymous and voluntary surveys that combined quantitative answers with open-ended text responses. We compared student responses based on gender, ethnicity, and school location [41, 42]; learned student views about whether SEFs should be competitive vs. non-competitive and optional vs required [43, 44]; and identified experiences and types of help that correlated with increased student interest in science and engineering [37, 45]. We believe that having this information helps educators develop best practices leading to more effective, inclusive, and equitable SEF learning opportunities, thereby enhancing successful student participation and outcomes.

Little attention has been given to the communication practices students employ during SEF participation even though students engage in these skills as part of authentic scientific activity, and *Obtaining, Evaluating, and Communicating Information* is one of the eight Science and Engineering Practices in the National Research Council framework [1, 2]. The goal of the current study was to provide a foundational analysis of the types of communication and presentation skills used by students in their SEF projects and the association of these skills with SEF outcomes. To accomplish this goal, we included the following question in our national surveys carried out during 2021-22 and 2022-23, “What types of communication and presentation skills did you use in your science fair project?” The possible answers were *literature review; research notebook; software to prepare tables, graphs, or images; written report; poster board preparation; PowerPoint presentation; and interview with the judges*. Poster board preparation and interview with the judges are skills frequently included explicitly as part of SEF scoring criteria [46]. The other skills about which we asked are elements of scientific inquiry from developing a research question to carrying out the experiments and collecting data to preparing the results for presentation to others. 1789 students answered the question. In this paper, we report and discuss the implications of their answers.

## Materials and methods

This study was approved by the UT Southwestern Medical Center (IRB #STU 072014-076). The study design entailed administering to students a voluntary and anonymous online survey using the REDCap survey and data management tool [47]. Survey recipients were U.S. high school students using Scienteer (www.scienteer.com) for online SEF registration, parental consent, and project management during the 2021/22, and 2022/23 school years. Although we treat the Scienteer SEF population as a national group of U.S. high school students, it should be recognized that these students come from seven U.S. states: Alabama, Louisiana, Maine, Missouri, Texas, Vermont, and Virginia. We have no information about the different locations where SEF fairs are held within each of the seven states.

After giving consent for their students to participate in SEFs, parents could consent for their students to take part in the SEF survey. However, to prevent any misunderstanding by parents or students about the possible impact of agreeing to participate in the survey, access to the surveys was not available to students until after they finished all their SEF competitions. When they initially registered for SEFs, students whose parents gave permission were told to log back in to Scienteer after completing the final SEF competition in which they participated. Those who did so were presented with a hyperlink to the SEF survey. Scienteer did not send reminder emails, and no incentives were offered for remembering to sign back in and participate in the survey. Since 2016, when we began surveying the national Scienteer cohort of SEF students, more than 4,000 students have completed surveys, an overall response rate of about 3%. Given that student participation in the surveys involved an indirect, single electronic invitation without incentive or follow-up, this level of response was not unexpected [48-50]. Nevertheless, one limitation of our findings is that the survey respondents may not fully represent the broader population of SEF participants. As in our previous work, we analyzed survey results for two consecutive years to increase consistency, reliability, and reproducibility. Our analyses are descriptive and exploratory in nature, intended to identify possible associations rather than demonstrate causal relationships.

The survey used for the current study can be found in supporting information (S1 Survey). The current version is similar to the original survey first adopted in 2015 [51]. However, since then new questions have been added about level of SEF competition; interest in a career in S&E; student ethnicity; school location (urban, suburban, rural); and reasons why or why not science fair experience increased the students’ interest in S&E. Beginning in 2021 we added the question *What types of communication and presentation skills did you use in your science fair project?* The possible answers were *literature review*; *research notebook*; *software to prepare tables, graphs, or images, written report*; *poster board preparation*; *PowerPoint presentation*; and *interview with the judges*. Students could select multiple skills independently. The students’ responses to this question are the focus of the current study.

In the figures, survey data are summarized as frequency counts and percentages. In some cases, associations between categorical variables were evaluated using chi-square tests of independence (p values calculated without Yates correction). No adjustments were made for multiple comparisons.

## Results

### Overview of survey responses about communication and presentation skills

Figure 1 shows the year-to-year comparison and the averages of students’ responses to the question about communication and presentation skills used in their SEF projects. Literature review was reported by the fewest students (15-20% across survey years). Poster board preparation was utilized by the greatest number (60-77%). The students’ responses were similar overall across the two survey years except for PowerPoint presentation, which showed greater variation than the other skills measured. In subsequent figures, we show the combined results for the two survey years.

**Figure 1.**
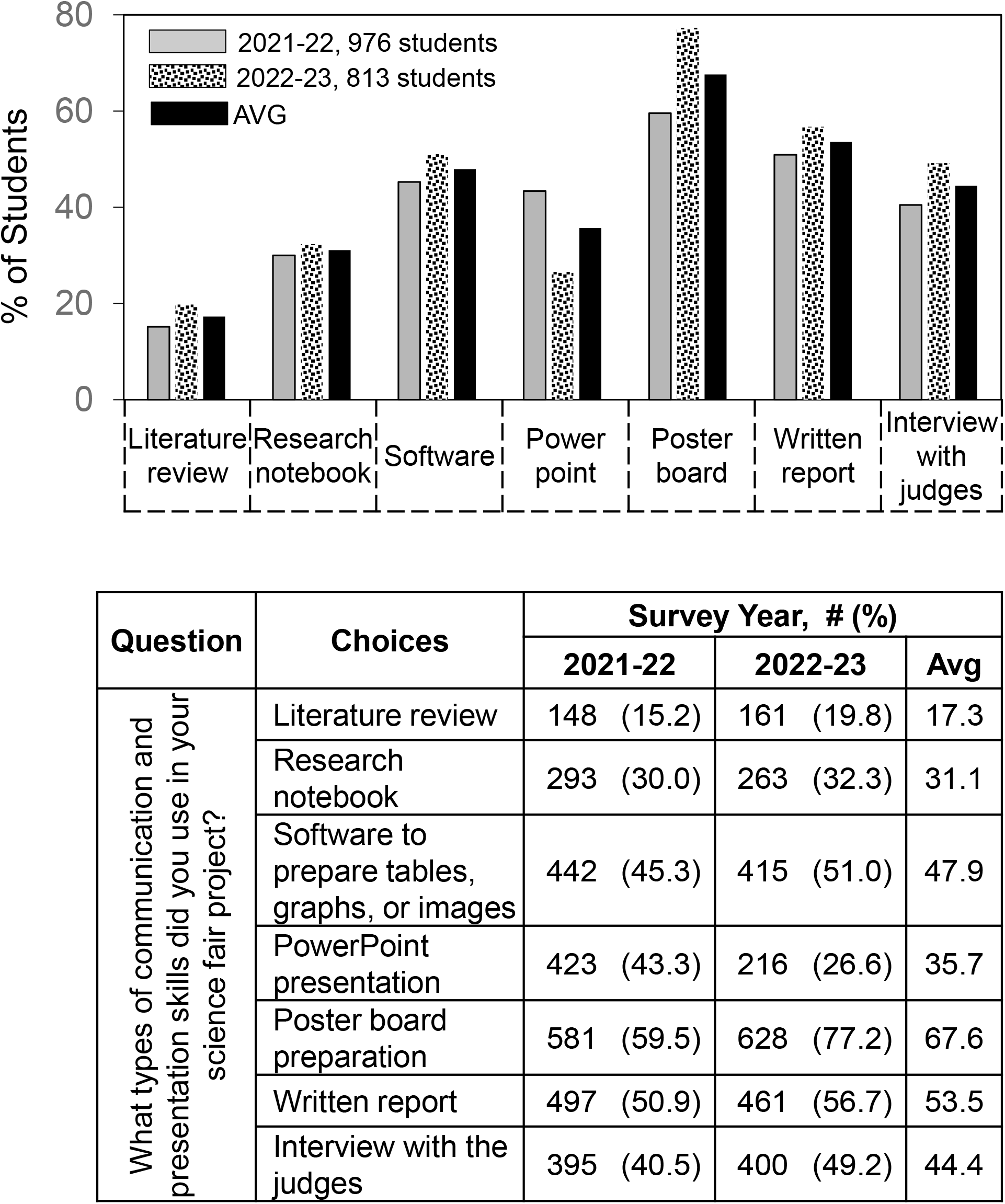
Year-to-year comparison and the averages of students’ responses to the question about communication and presentation skills used in their SEF projects.

Figure 2 shows the overall extent of communication and presentation skill use. The number of skills used ranged from zero (4.1% of the students) to all seven (2.5% of the students). On average, the mean number of skills used per student was 2.99. Subsequent figures show how individual skills differed according to student demographics, help received, and SEF outcomes. These analyses identify associations between communication practices and outcomes but cannot determine causality.

**Figure 2.**
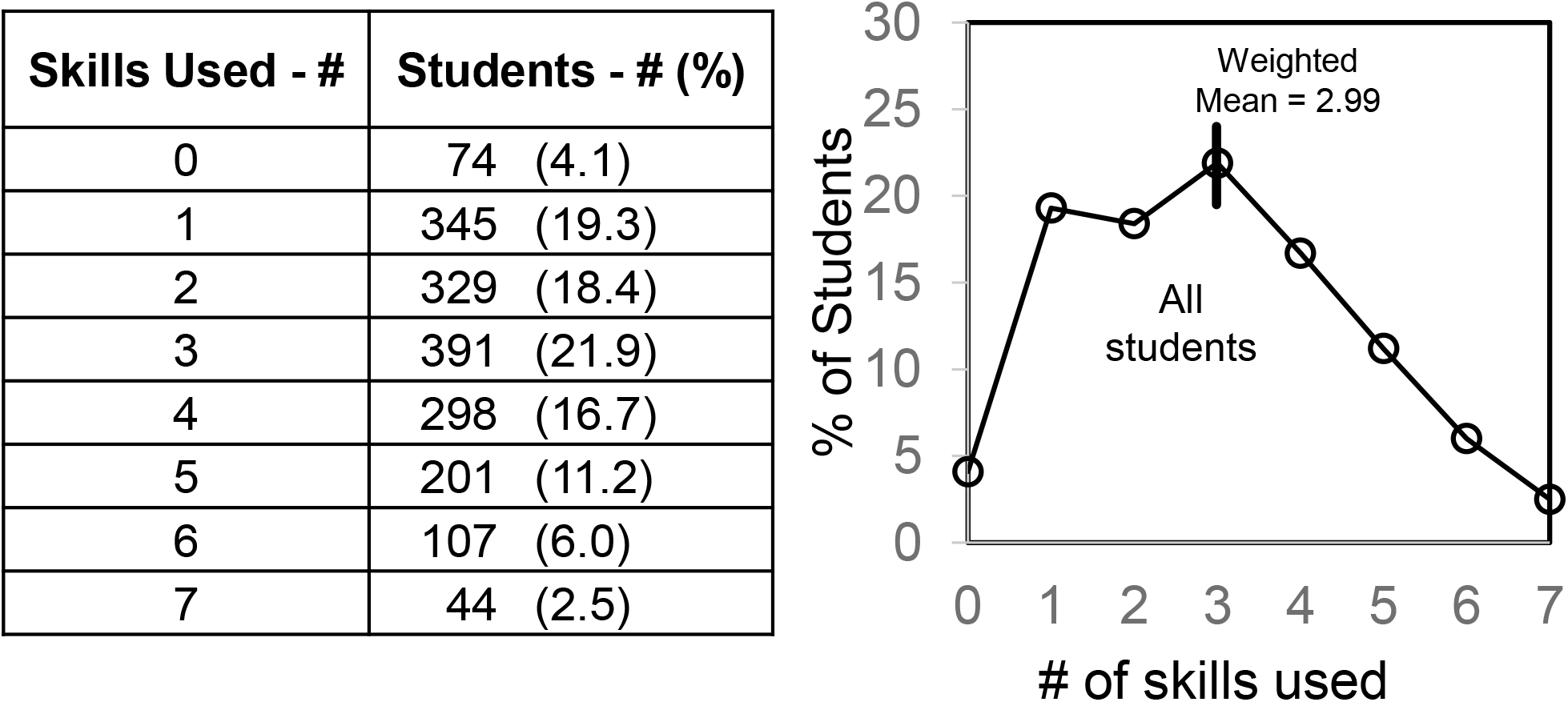
Overall frequency of skill use.

### Demographics

In Figures 3 and 4, survey results were analyzed to determine if differences in particular skill use were associated with gender or ethnicity. Figure 3 shows survey responses based on gender. Overall, use of communication and presentation skills showed only small gender differences in the various categories. Females reported slightly higher percentages in most categories.

**Figure 3.**
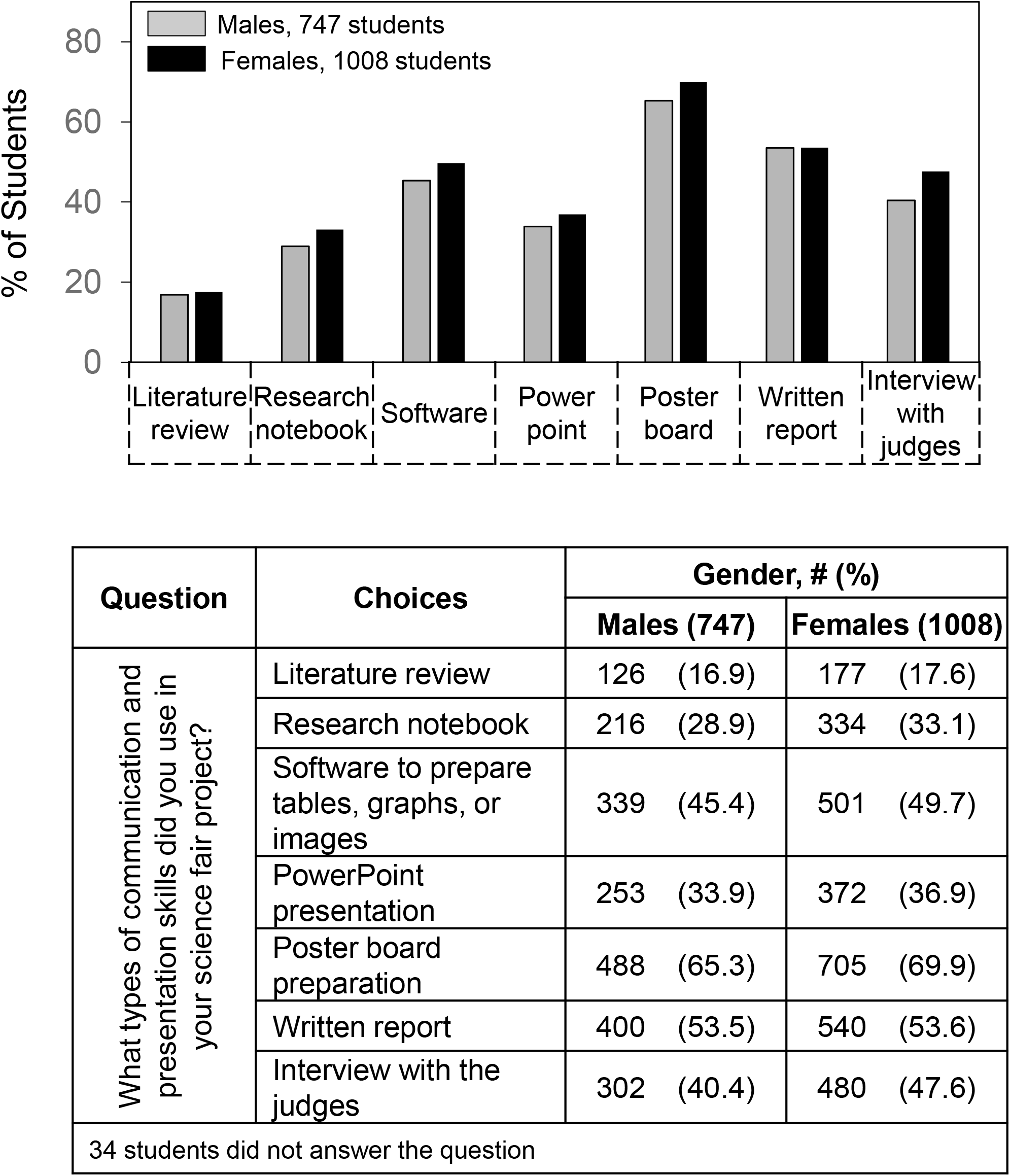
Skill use and gender.

**Figure 4.**
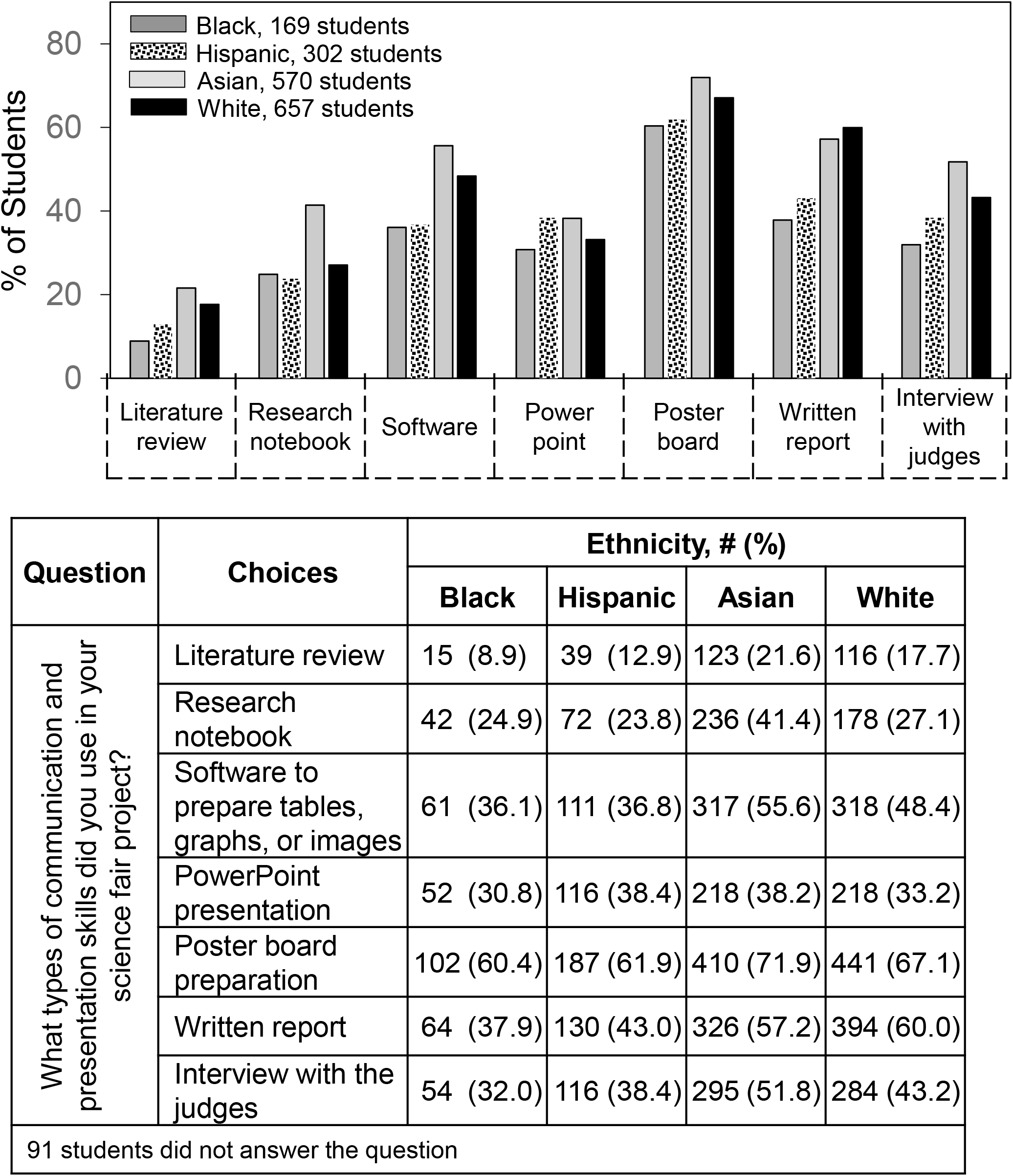
Skill use and ethnicity.

Figure 4 shows survey responses based on ethnicity. Unlike gender, greater variation was observed across ethnic groups. Use of PowerPoint presentation and poster board preparation were similar for all groups. Asian students reported higher percentages of skill use per student (3.4); White students were intermediate (3.0); and Hispanic and Black students were lower (2.6 and 2.3). The most notable differences were Asian students for higher use of literature review, research notebook and interview with the judges; and Asian and White students for higher use of software and written report.

### Help students received

In Figures 5 and 6, survey results were analyzed to determine if differences in particular skill use were associated with types of help that students received. Figure 5 shows survey responses based on help from parents (alone), teachers (could also include parents), and scientists (could also include parents and teachers). Except for use of PowerPoint presentation, students receiving help from scientists were more likely to report a higher percentage of skill use followed by students receiving help from teachers and then from parents. In the case of use of a research notebook, students with help from scientists were twice as likely (60% vs. 30%) to report use of the skill.

**Figure 5.**
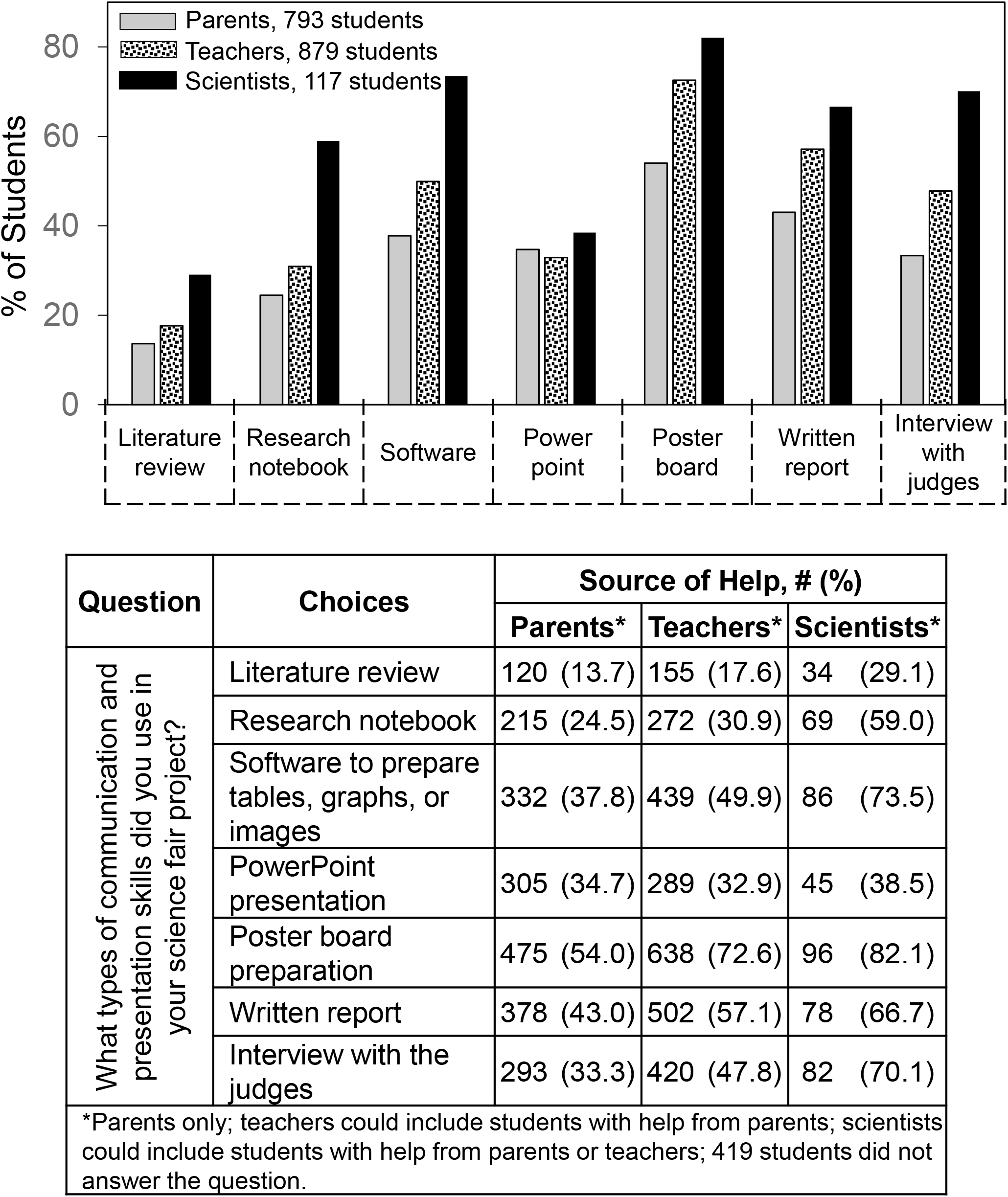
Skill use and help from parents, teachers, and scientists.

**Figure 6.**
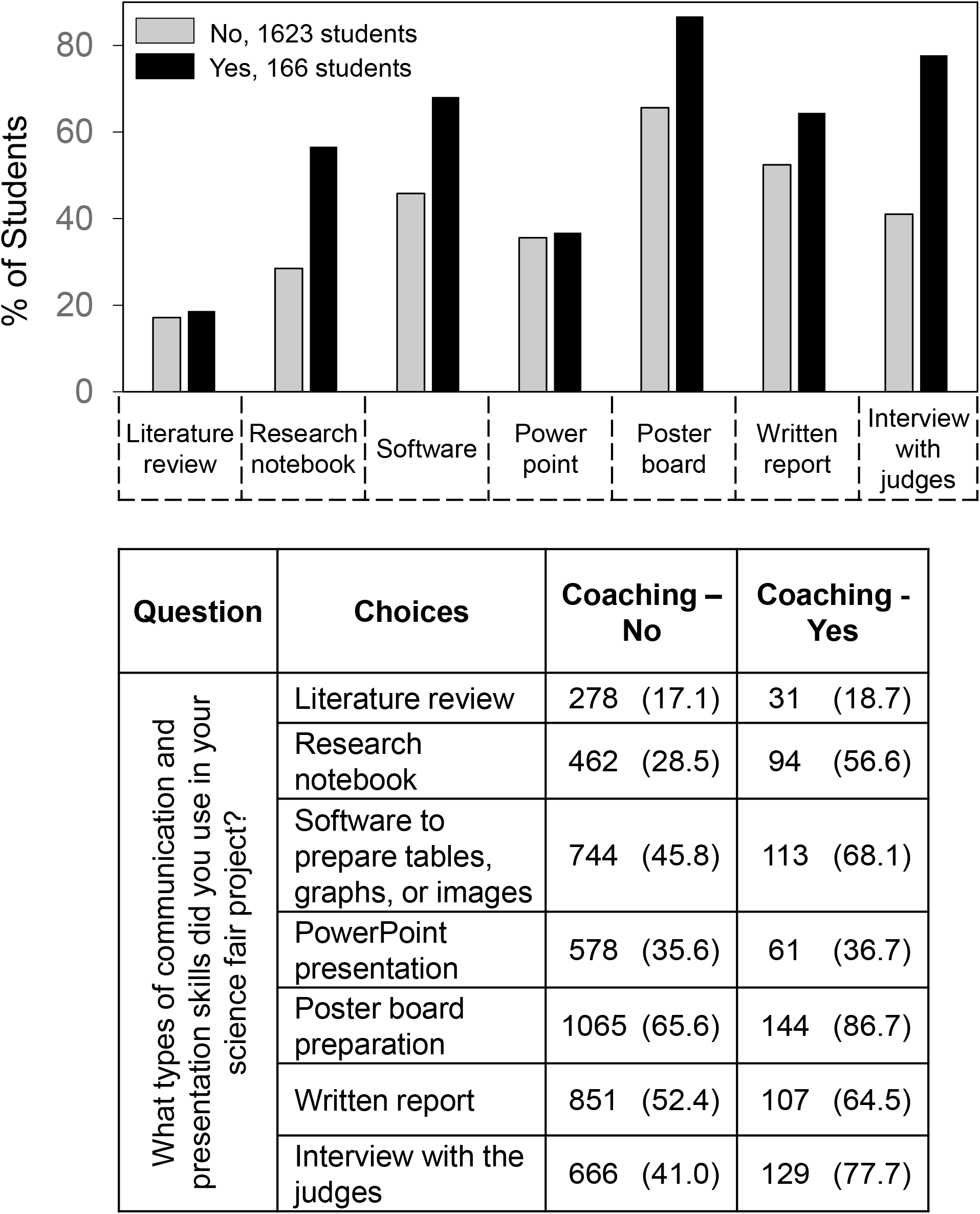
Skill use and coaching for the interview.

Figure 6 shows survey responses based on whether students received help coaching for the interview. Except for literature review and use of PowerPoint presentation, help coaching for the interview was associated with an increase in skill use. As in the case of help from scientists, the largest difference was in reported use of a research notebook (57% vs. 29%). Also, students who received coaching for the interview reported higher likelihood of interviews with judges (78% vs. 41%).

### SEF outcomes and skill use

Figure 7 shows associations between skill use and progressively higher levels of SEF competition. In this comparison, students could participate in SEF competitions ranging from school to state level. The results shown are for the highest level of SEF competition in which students reported competing. The average number of skills used by students increased with higher levels of competition - school (2.6), district (3.5), regional (3.9) and state (4.4). Students in the school only group reported especially lower percentages of use of research notebooks and interview with the judges compared to students in the other competition levels, whereas use of written reports was similar across competition levels.

**Figure 7.**
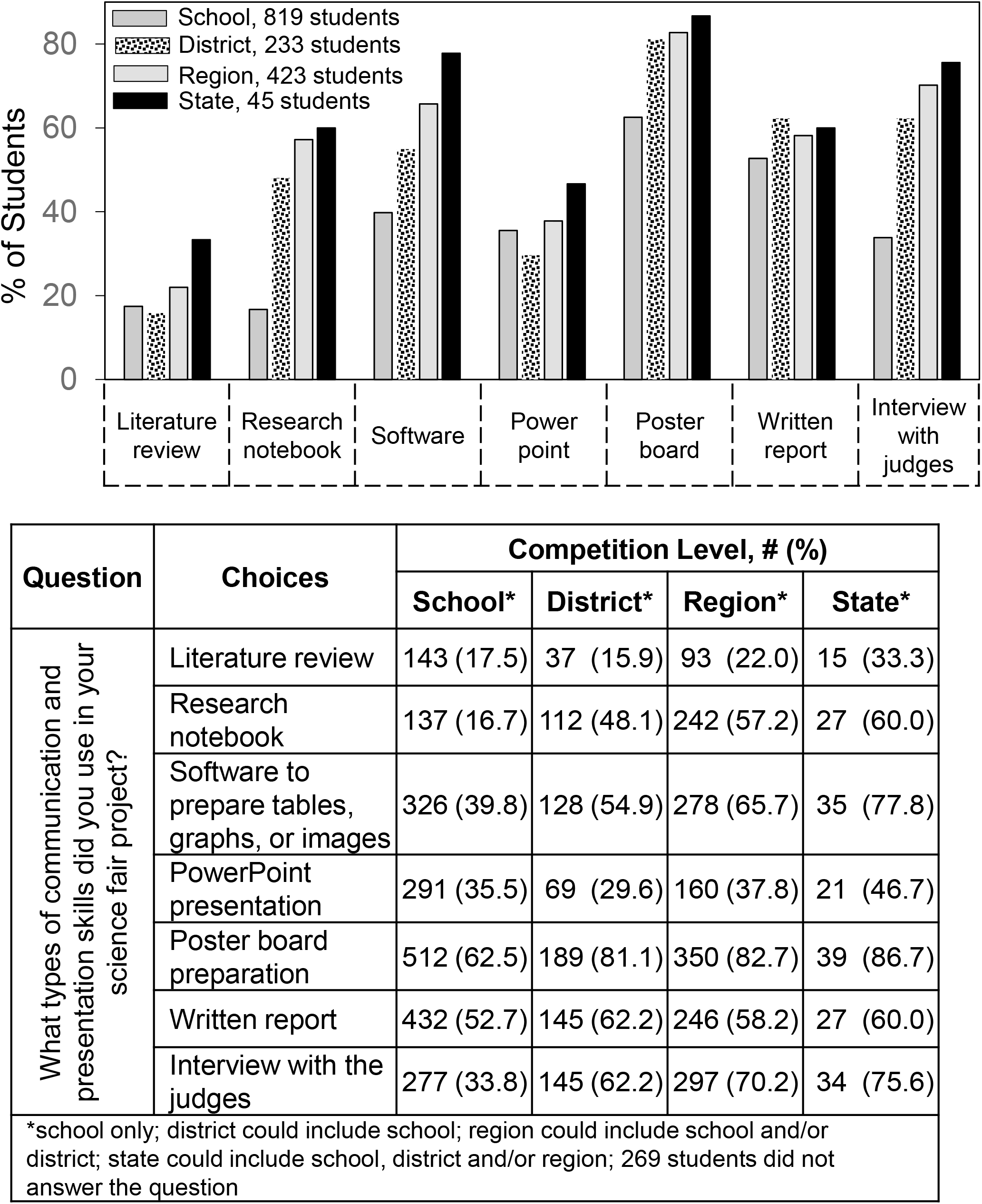
Skill use level of SEF competition.

Figure 8 shows use of communication and presentation skills in relation to whether students were interested in a career in S&E. The overall trend was for students who selected having a career interest to report greater skill use compared to students with unsure or no interest in an S&E career. Four of the skills about which we asked students showed a strong association (*p*<.001) with greater interest – research notebook; software to prepare tables, graphs and images; poster board preparation; and interview with the judges. Of this group, use of a research notebook stood out more than three times as likely to be used by students interested in an S&E career.

**Figure 8.**
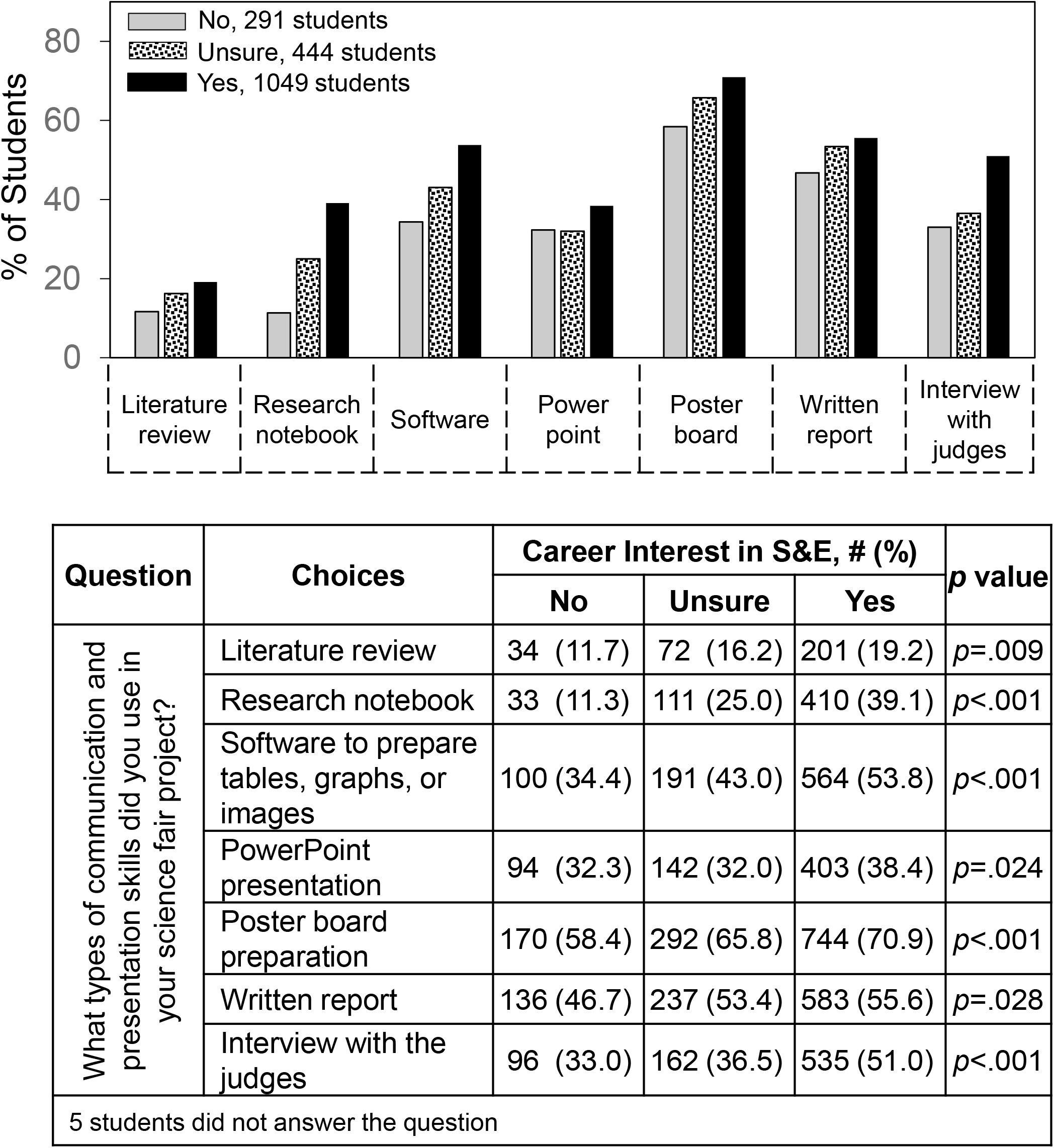
Skill use and student interest in an S&E career.

Figure 9 shows use of communication and presentation skills in relation to whether students indicated that participation in SEFs increased their interest in S&E. Similarly to the results regarding interest in an S&E career, four of the skills about which we asked students showed a strong association (*p*<.001) with increased interest – research notebook; software to prepare tables, graphs and images; poster board preparation; and interview with the judges. Use of a research notebook and interview with judges stood out and were twice as likely to be used by students who indicated that SEF participation increased their interest in S&E.

**Figure 9.**
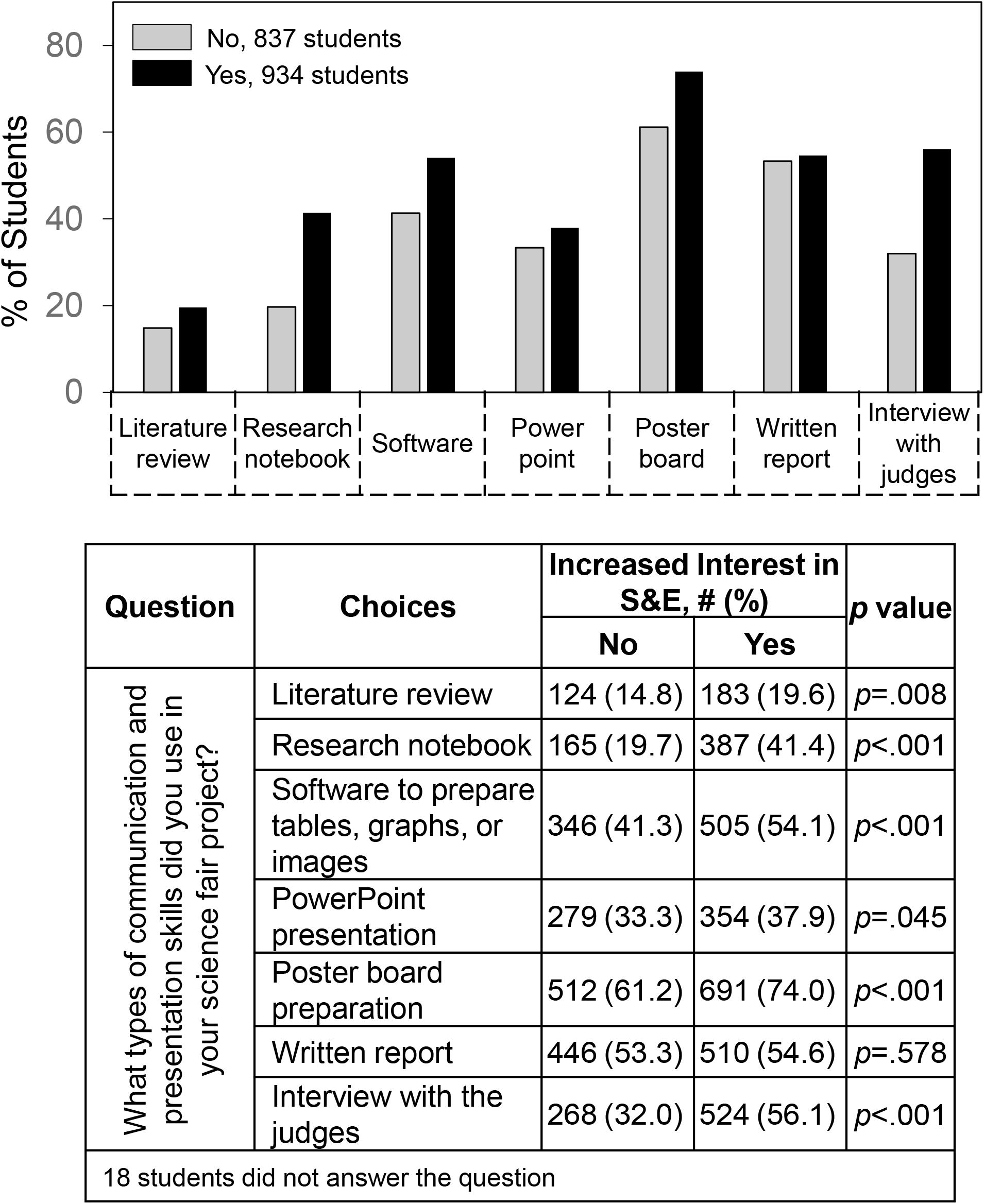
Skill use and whether SEF participation increased student interest in S&E

## Discussion

Communication and presentation skills are integral parts of scientific inquiry and involved in choosing research questions, documenting evidence, interpreting findings, and reporting conclusions. In this paper, we present findings based on quantitative answers by 1789 students who participated in our SEF surveys during 2021-22 and 2022-23 and answered the question “What types of communication and presentation skills did you use in your science fair project?” The possible answers were *literature review; research notebook; software to prepare tables, graphs, or images; written report; poster board preparation; PowerPoint presentation; and interview with the judges*.

The possible answers connect with different phases of scientific inquiry. Literature review and research notebook are part of developing research questions and data collection. Software to prepare tables, graphs, or images, and PowerPoint presentation contribute to analyzing research and developing a presentation. Written reports, poster board preparation, and interviews with the judges provide the opportunity to present the findings. Among these, only poster board preparation and interview with the judges are skills explicitly included as part of typical SEF scoring criteria. Judges are advised to examine the student notebook but not to score it [46].

On average, students indicated they used three skills, which most often were those included in SEF project scoring -- poster board preparation and interview with the judges. These two options were selected more than twice as frequently as literature review and research notebook although the latter two options are implicit in scientific inquiry. As SEFs are now scored, the latter skills can only be assessed indirectly through inclusion in sections of the poster board or written report or as part of the interview. If SEF scoring was modified to include literature review and research notebook explicitly, then students might be encouraged to give these skills more direct attention. Currently, presentation of the results appears to get the most attention from students.

Demographic studies showed few gender differences between students. However, we observed ethnicity-dependent differences. Asian students reported a higher percentage of skill use in most categories. Overall skill use per student was highest for Asian students (3.4) followed by White students (3.0) and then Hispanic and Black students (2.6 and 2.3). Since the differences between ethnic groups were evident for interview with the judges but not poster board preparation, one interpretation of this finding is that students in all groups had the opportunity to present their work by poster even if they didn’t get an interview with judges.

Previously we found that Asian and Hispanic students indicated more so than Black and White students that SEF participation increased their interests in S&E [41, 42] and made more free text positive comments about their SEF experiences [37]. The present study suggests that differences in the students’ uses of communication and presentation skills cannot account for the differences in the students’ attitudes towards SEF experience that we reported previously.

More than 50% of students receive help from parents and teachers. Less than 10% receive help from scientists. However, compared to other students, those that received help from scientists were much more likely to report use of a literature review, research notebook, and software, all typical features of scientific inquiry. Help from teachers also had an impact but mostly on the skills involved in presentation of the research rather than inquiry itself. Students who received coaching for the interview also indicated greater use of a research notebook and software. These differences provide additional support for the conclusion that help from scientists and coaching should be expanded as much as possible to advise students about their SEF projects [32-34, 41, 52].

The finding that the average number of skills used by students increased with higher levels of competition - school (2.6), district (3.5), regional (3.9) and state (4.4) – is consistent with the idea that the use of skills we are measuring is an indication of successful student engagement with scientific inquiry. The trend was not just overall but for almost every type of skill except written report. However, these differences might reflect not only the students’ abilities but also in part their school’s involvement. That is, not all schools or school districts have the financial and other resources to make it possible for students to participate in SEFs beyond the school only level. The findings suggest that for school only SEFs, use of written reports may provide an opportunity for students to present their SEF research even if an interview to verbally present the work with judges is unavailable.

Whether or not students indicated that they are interested in a career in S&E and whether SEF participation increased their interest in S&E are key indicators of successful SEF outcomes. Four of the skills about which we asked students showed a strong association (*p*<.001) with successful outcomes – research notebook; software to prepare tables, graphs and images; poster board preparation; and interview with the judges. Of this group, use of a research notebook stood out, more than three times as likely to be used by students interested in an S&E career and twice as likely to be used by students who indicated that SEF participation increased their interest in S&E. Ironically, even though students do not select research notebooks as one of the more frequently used skills in SEFs and are not scored on this skill explicitly, the science education literature has long emphasized the value of research notebooks to help students engage in reflection, interpretation, and conceptual understanding of their work [53-56] as well as improving literacy overall [57-59].

Several limitations of our study are worth noting. One is that we treat the Scienteer SEF population as a national group. However, it should be recognized that these students may not be truly representative of a national sample since they come from only 7 U.S. states and only attend high schools where SEFs are available. In addition, we cannot be sure that the 3% response rate of survey respondents is representative of the overall high school student population participating in SEFs. Also, year-to-year student answers regarding PowerPoint showed more variation than other skills. Finally, we cannot be sure that students mean the same thing when they choose the different skills.

In conclusion, our findings help understand the ways in which communication and presentation practices were associated with SEF participation and outcomes and how these practices can be encouraged in different ways by engagement with teachers, scientists, and coaching. Surprisingly, inquiry practices of literature review and research notebook, which provide the underlying framework for developing and carrying out scientific inquiry, are least selected by students, perhaps because of how SEFs are scored. More than any other skill, use of research notebooks was associated with positive SEF outcomes.

## Acknowledgments

FG was supported by the UTSW Robert McLemore Professorship. Use of the REDCap survey and data management tool was facilitated by the UTSW Department of Information Resources and Clinical and Translational Science Training Program, NIH grant UL1TR001105. The REDCap funders had no role in study design, data collection and analysis, decisions to publish, or preparation of the manuscript. We are grateful to Russell Cowen and Rocky Slavin who were managers of Scienteer Technologies at the time we carried out our studies and incorporated the parental consent and SEF survey REDCap links into the Scienteer website.

## Supporting information

S1 Survey. Quantitative student responses to the SEF survey year by year.

